# The Effects of Flow Speeds on Smooth Pursuit Tracking and Active Sensing Movements of Weakly Electric Fish

**DOI:** 10.1101/2023.11.02.565374

**Authors:** Emin Yusuf Aydin, Burcu Unlutabak, Ismail Uyanik

## Abstract

Weakly electric fish employ refuge-tracking behavior to survive, seeking and utilizing hiding places to shield themselves from predators and unfavorable environmental conditions. This adaptive mechanism enables them to minimize the risk of predation, maintain optimal electrocommunication, and adapt to changing surroundings. While studies have explored smooth pursuit tracking and active sensing movements of these fish in stationary environments, limited emphasis has been given to how varying flow speeds in their natural habitats may impact these behaviors. This study addresses this gap by investigating the effects of different flow speeds on smooth pursuit tracking and active sensing movements in weakly electric fish. Active sensing provides sensory data and multisensory integration processes and combines this data to create a holistic perception of the environment. The synergy between these processes is fundamental for enhancing an organism’s sensory capabilities and enabling it to adapt and interact effectively with its surroundings. For this study, a specialized experimental setup was designed and built to facilitate refuge-tracking behavior under controlled flow conditions. The experiments involved *Apteronotus albifrons* fish exposed to visual and complex electrosensory stimuli, which consisted of a sum of sine signals. Data was recorded for different sensory conditions, including variations in flow speeds, illumination levels, and refuge structures. The analysis revealed that increased flow speeds correlated with reduced tracking gain and phase lag in the fish. Additionally, it was observed that active sensing movements were more pronounced in dark conditions. These findings highlight the significant impact of flow speeds on smooth pursuit tracking and active sensing movements and emphasize the importance of studying these behaviors within the context of water flow. Understanding the biological motivations underlying these effects is vital for their potential application in engineering fields.

## 1. INTRODUCTION

Weakly electric fish have long been the focus of scientific research due to their remarkable ability to produce and sense electric fields (Comertler and Uyanik, 2021; Gabbiani et al., 1996; Metzen et al., 2016; Von der Emde, 1999). They have a complex sensory system that integrates several senses, including vision and electrosensing. Specifically, weakly electric fish have an impressive electrosensing system. These fish use specific electric organs to generate weak electric fields, and their electroreceptor organs allow them to detect changes in these fields. Thanks to this electrosensing system, they can explore their surroundings, find prey, and communicate with other conspecifics (Ammari et al., 2014; Biswas et al., 2018; Gabbiani et al., 1996; Heiligenberg and Bastian, 1984; Metzen et al., 2016). Moreover, weakly electric fish have vision systems suited to their unique habitat. They can recognize visual cues in low light, which helps with prey detection, object recognition, and social interactions. Thus, these fish can develop their senses and exhibit the proper behavioral responses by integrating electrosensing and visual data at different levels of cerebral processing (Bastian, 1982; Gottwald et al., 2018; Kareklas et al., 2017; Moller, 2002; von der Emde, 2004).

The species *Apteronotus albifrons* (Linnaeus,1766), known as a weakly electric fish, exhibits fascinating changes in its behavior while tracking a refuge, depending on the availability of different sensory cues. Refuge tracking refers to the fish’s behavior of closely following the movement of a refuge by swimming forward and backward to remain within its confines. This behavior has been observed in natural surroundings, such as when fish seek refuge in fallen palm trees or other vegetation, as well as in controlled laboratory conditions using PLA (Polylactic acid) tubes or similar refuges. The fish adapt their reliance on visual and electrosensory cues throughout the tracking process, depending on the significance/salience of each cue, with electrosensory cues becoming particularly crucial in complete darkness. While fish primarily show smooth and linear tracking movements, they also exhibit a type of fore-aft movement that is not directly connected to the refuge’s motion (Uyanik et al., 2019).

Using these active movements for exploring and examining their environment, the fish gather specific sensory data that help them make better decisions and increase their chances of survival. Active sensing refers to the intentional actions of animals to actively expend energy to gather information from their environment (Nelson and MacIver, 2006). To this end, animals use sensory systems such as vision, hearing, smell, touch, or electrosensing, and they engage in active sensing for various reasons related to their survival and adaptation to their environment. Active sensing can improve an animal’s perception by actively manipulating the environment to enhance the quality or usability of this sensory information. This sensing enables animals to obtain crucial environmental information, such as locating food sources, detecting predators, identifying potential mates, or finding suitable habitats (Stamper et al., 2012). For example, bats emit ultrasonic sounds and listen for echoes to locate their prey in dark environments. This active echolocation allows them to perceive their environment in greater detail than passive perception alone (Jones et al., 2021).

The interest in studies investigating the mechanisms of active sensing behavior is increasing due to biological implications and engineering applications. However, studies conducted so far with weakly electric fish solely focus on fish’s response in stationary environments. Many species of weakly electric fish live in flowing water bodies such as rivers and streams (Winemiller and Adite, 1997). The flow rate of the river affects various aspects of the fish’s life, including navigation, foraging, and communication. Weakly electric fish need to adapt to different flow velocities to find food, avoid predators, and move efficiently through their environment. This important factor has generally been overlooked in many studies in the literature, and they have been carried out in stationary waters (Stamper et al., 2010; Tan et al., 2005). Since these fish experience varying flow speeds that possibly affect their active sensing movements in their natural habitats, it is important to account for this factor to gain a better understanding of fish’s behavior. This study addresses the gap in the literature by presenting a novel investigation of the effects of different flow speeds on the smooth pursuit tracking and active sensing movements of weakly electric fish.

## 2. MATERIALS AND METHODS

Five individual adult *Apteronotus albifrons*, aged 18-24 months, were used to conduct refuge-tracking experiments. Experiments were carried out with the permission of the Hacettepe University Animal Experiments Ethics Committee (No: 2023/05-07). During the experiments, the water temperatures and pH values were maintained at 25 ± 1°C and 7.2, respectively. Fish were fed with frozen bloodworms once a day. The care of the fish and the experimental procedures were carried out in accordance with ethical rules, minimizing animal stress. The fish used in the experiments were placed in the experimental setup without distinguishing them as male or female. In the process of this study, no fish were killed. Before the experiments started, the fish were kept in the experimental setup for two hours to acclimatize to the experimental environment.

### 2.1. Experimental Apparatus

We designed and built a unique experimental setup to carry out the experiments in this study. This setup can be thought of as a specialized aquarium system in which the fish perform their tracking behavior in a moving PLA refuge. In this experimental setup, the refuge movements are provided by a high-precision linear DC motor (Maxon motor 380795, Maxon Group, Switzerland) that can move in a single axis. The bottom part of the fish measuring 25 cm x 50 cm under the test area was cut off and replaced with glass. In this case, it was ensured that the camera images of the fish were recorded from the bottom. The behavioral response of the fish to the refuge movements was recorded by a Near Infrared Camera (Basler ace acA1300-60gm-NIR, Basler AG, Lübeck, Germany) placed under the experimental setup. The flow speeds of the water are provided by T200 thruster (Blue Robotics Inc., Torrance, CA) placed in the setup. One critical point was obtaining a water flow as constant as possible at every moment of the assembly. In this context, perforated mechanical filters (honeycomb) were used to regulate the water flow at the connection point where the water inlet is attached to the experimental setup. The system’s general electronic and software architecture runs on the Robot Operating System (ROS) Melodic. The real-time online loop frequency of the system was determined as 25 Hz. The leading software of the system works via the Jetson Xavier NX (Nvidia, Santa Clara, CA) card using the ROS Melodic. The main tasks of this board were (1) starting and controlling the relevant experiment sequence according to the commands from the interface, (2) transmitting the necessary motion commands to the motor driver and processing the feedback about the motor position and fish positions. The collection of squares can sort it. Motion control of the linear motor is provided by an EPOS 4 50/5 motor driver (Maxon Group, Switzerland) to be driven via the Jetson Xavier NX. Single-axis motion control of PLA refuge is provided with millimeter precision.

### 2.2. Experiment Procedure

Experiments were performed in light (∽300 lux) and dark (∽0.04 lux) conditions with low conductivity (∽40 mS/cm) conditions. PLA refuge, 14 cm in length and 4 cm wide, with windows, was used in the experiments. The experiments were carried out at four different flow rates: stationary (0 cm/s), low (4.5 cm/s), medium (11 cm/s), and high (15.5 cm/s). These rates were calculated by the transit time of a ping pong ball through a linear tube system. For each speed value used in these experiments, we had 20 experiment video records. We used the time that the ball passed through the 30 cm tube from the videos and thus reached these rates. The refuge was moved with a single sinusoidal input as the sum of 13 sinusoids at different frequencies (0.10, 0.15, 0.25, 0.35, 0.55, 0.65, 0.85, 0.95, 1.15, 1.45, 1.55, 1.85, 2.05 Hz). Experiments for each fish were repeated five times, carried out from low to high speed, with each experiment for sixty seconds (1500 frames). We performed a total of 500 trials for the current study. The data for each fish was collected over 1-2 weeks. As we have observed in previous studies, fish did not show long-term adaptation or changes in tracking performance over time (Cowan and Fortune, 2007; Stamper et al., 2012; Uyanik et al., 2020).

### 2.3. Data Analysis

Using our custom-built code written using Python 3.9, the fore-aft position of the fish was tracked from the recorded video. Fish and refuge positions were digitized using our custom image processing code implemented in MATLAB 2023b (MathWorks, Natick, MA). For each trial, we measured the trajectory of the refuge, *r(t)* and the fish, *y(t)*. Fish position was measured using a custom template-based video tracking algorithm centered on the black and white difference used just at the end of the fish head (Fig.2).

**Fig 1.**
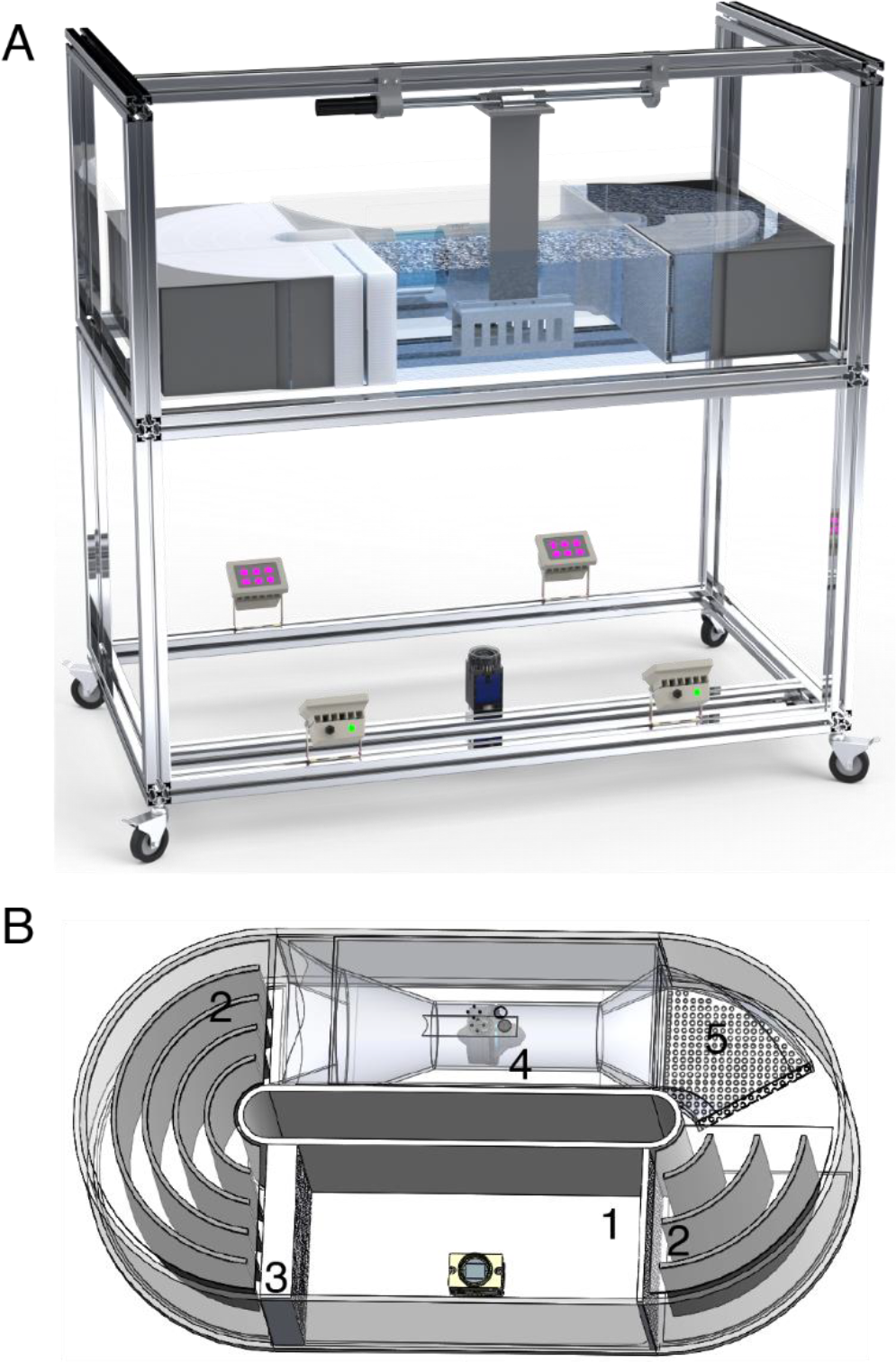
Experimental apparatus and setup. **A)** All elements in the experimental setup (Bottom: IR LEDs and camera, Middle: experimental setup, Up: Motor and linear actuator system **B)** Experimental setup of the flow tank with an aerial view. The camera was placed at the bottom of the experiment section (1) of the flow tank. There are breakwaters (2) and honeycombs (3) on both sides of the experiment area for a more accurate flow of water. A thruster (4) absorbs water through an absorber (5) and pumps toward the test area.

**Fig 2.**
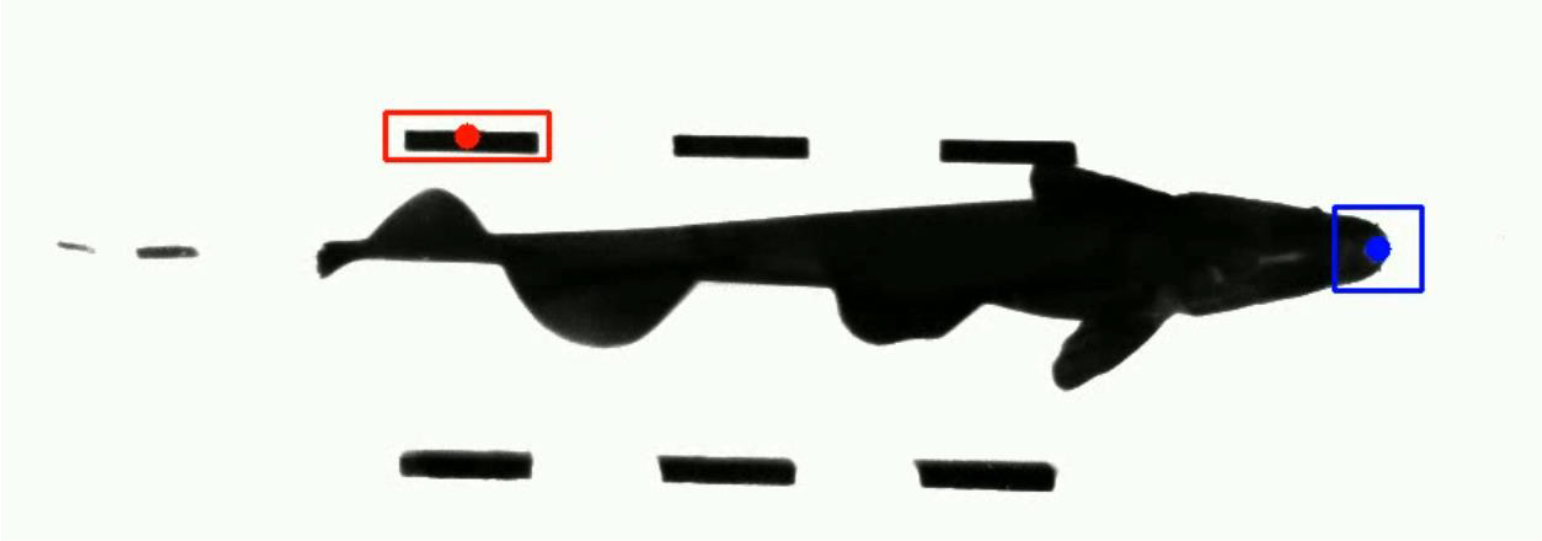
A screenshot from custom-built template-based video tracking. The blue dot marks the position of the fish, *y(t)* and the red dot corresponds to the position of the refuge, *r(t)*.

The Discrete Fourier Transform (DFT) represents the time domain signals *r(t)* and *y(t)* as complex-valued functions of frequency, *R*[ω] and *Y*[ω]. These complex numbers can also be represented in polar coordinates in terms of their magnitude, |*Y*[ω]|, and phase ∠*Y*[ω]. For the sum of sines wave input trajectories, the DFT of *R*[ω] is represented as discrete spikes at the refuge frequencies and zero at all other frequencies. In contrast, the DFT of the fish movement *Y*[ω], typically as power over a broader range of frequencies (Uyanik et al., 2019) (Fig.3).

**Fig 3.**
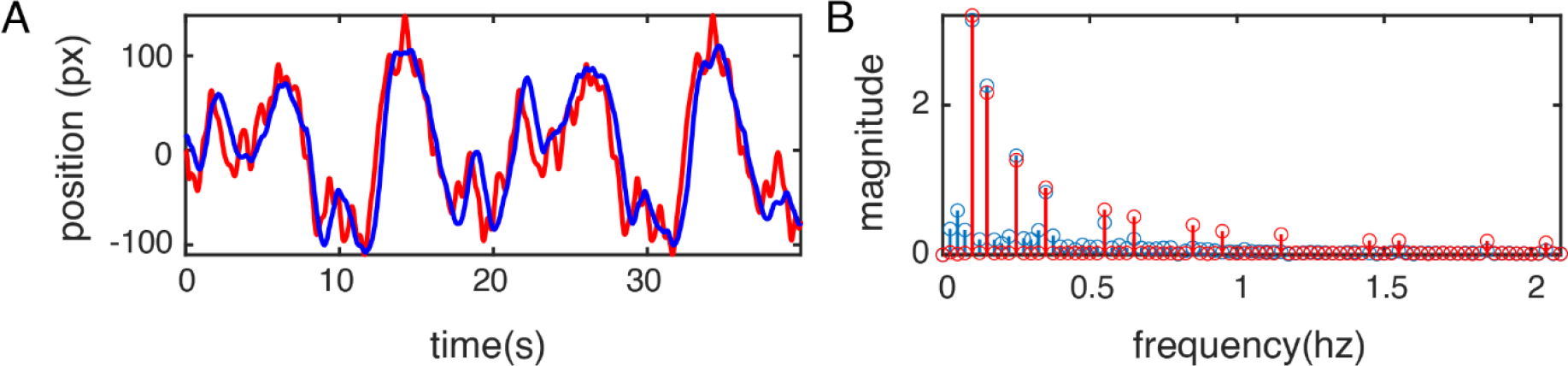
An example tracking result. Left: tracking result in time domain and Right: DFT of the tracking result in frequency domain. The blue line represents the fish movements, *y*(*t*), and the red line represents the refuge movements, *r*(*t*).

Frequency-response plots describe the response of a system by comparing the output signal, *Y*[ω], to the input signal, *R*[ω],using two measures, gain and phase. For each frequency ω_0_, the gain is calculated as the ratio of the signal magnitudes, |*Y*[ω_0]_]|/|*R*[ω_0_]|, and phase is computed as the difference between signal phases, ∠*Y*[ω_0_] − ∠*R*[ω_0_]. The frequency-response plot is only evaluated at the stimulus frequency, as the gain ratio and phase lag are not defined where the stimulus magnitude is zero, i.e.,, *R*[ω] = 0 (Uyanik et al., 2019).

### 2.4. Statistical Analysis

To examine the effects of different sensory conditions on weakly electric fish’s tracking behavior, we conducted a repeated measures ANOVA using within-subject factors of illumination (2: light and dark), refuge structure (2: window and no window), and flow speed (4: stationary, low, medium, high) (Jamovi, version 2.3.28, www.jamovi.org). Our outcome measures included smooth pursuit tracking and active sensing movements of weakly electric fish as indexed by error values/metrics which are root mean square (RMS) for time domain and sum of weighted frequencies for frequency domain analysis. We calculated the mean error values across experimental trials and conducted the analysis on the mean values. Statistical significance for all statistical tests was set at p≤0.05. All results are reported as means ± standard deviations. For pairwise comparisons and for follow-up of significant interactions, we used Bonferroni correction. We checked the normality of data visually using Q-Q plots. We also checked for sphericity assumption using Mauchly’s test of sphericity and we used Greenhouse-Geisser correction due to sphericity violations.

## 3. RESULTS

We examined the effects of the following sensory conditions: 1) different flow speeds, 2) illumination and 3) refuge structure on the smooth pursuit tracking and active sensing movements of weakly electric fish. Here we report the results of our experimental trials evaluating the impact of these factors on the fish’s tracking performance. First, we present the results about smooth pursuit tracking performance (section 3.1). Second, we present the results about active sensing movements (section 3.2).

### 3.1. The effects of illumination, refuge structure, and flow speed on smooth pursuit tracking

The smooth pursuit tracking performance of weakly electric fish was measured using two distinct approaches: the time domain and the frequency domain. First, we examined the effects of testing conditions of the time domain. The difference between the reference entity (refuge) and the experimental subject (fish) was calculated using the RMS parameter. We found a main effect of illumination, *F* (1,23) = 53.095, *p* < .001, *η*_*p*_^*2*^*=* 0.698 indicating better tracking performance in light condition (*M* =54.20, *SD* =17.50) than dark condition (*M* =80.90, *SD* =16.80). The analysis showed that refuge structure had a significant effect on the tracking performance, *F* (1,23) = 8.426, *p* = 0.008, *η*_*p*_^*2*^*=* 0.268. Contrary to our expectations based on the literature, the tracking performance of the fish was better in no window condition (*M* =66.80, *SD* =16.80) than in window condition (*M* =68.40, *SD* =24.20). Finally, there was a main effect of flow speed on the tracking performance, *F* (3,69) = 16.512, *p* < .001, *η*_*p*_^*2*^*=* 0.418. The fish’s tracking performance was the best in stationary (*M* =55.30, *SD* =14.00) compared to other speed levels, *p* < .001. However, there was no significant difference across low (*M* =70.70, *SD* =19.90), medium (*M* =72.40, *SD* =20.10), or high levels (*M* =71.20, *SD* =14.30) (Fig.4).

**Fig.4.**
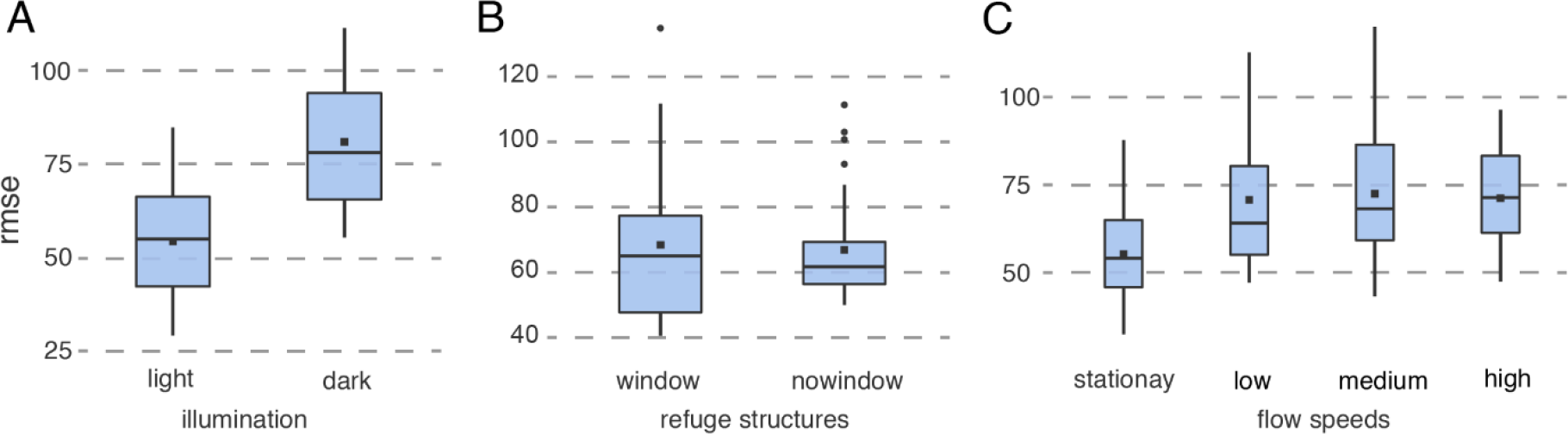
Smooth pursuit tracking across various sensory conditions in the time domain

In addition to the main effects, we found a significant interaction between illumination and flow speeds, *F* (3,69) = 3.323, *p* < .05, *η*_*p*_^*2*^*=* 0.126. Post hoc comparisons using Bonferroni correction showed that tracking performance was worse in light window condition than in light no window condition, p < .05. On the other hand, tracking performance was better in light window condition than in dark window condition. Moreover, tracking performance was better in light no window condition than dark window and dark no window conditions ps < .001.

Next, we examined the tracking parameters in the frequency domain. For this, the responses to 13 frequencies given as input were used. Low frequencies were prioritized in this metric, which was created using a weighting system. This system works using the following equation:

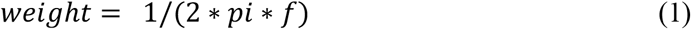

where *f* is a given frequency. Therefore, as *f* increases, the weight decreases, that is, the effect of that frequency on the result decreases. We divided the total result from this weighting by the number of frequencies (13) and defined it as the average error corresponding to a single frequency, which is frequency domain tracking error (FTE). Additionally, since it is difficult and ethically unsuitable to conduct experiments on fish, we applied the bootstrap method for the frequency domain. With this method, the long data obtained from the fish were randomly divided into 10 parts via custom MATLAB cade and analyzes were made with these parts.

As a result, we observed a significant impact of illumination conditions on tracking performance, with a noteworthy difference (*F* (1,239) = 535.391, *p* < .001, *η*_*p*_^*2*^*=* 0.691). It revealed that tracking performance was superior under ligt condition (*M* =0.297, *SD* =0.0933) compared to dark condition (*M* =0.392, *SD* =0.103). Our analysis also indicated that the design of the refuge structure had a notable influence on tracking performance (*F* (1,239) = 94.436, *p* < .001, *η*_*p*_^*2*^*=* 0.283). Surprisingly, contrary to our expectations based on existing literature, tracking performance was better in conditions with no window (*M* =0.325, *SD* =0.0933) than in conditions with windows (*M* =0.365, *SD* =0.117). Furthermore, we observed a main effect of flow speed on tracking performance, *F* (3,717) = 36.159, *p* < .001, *η*_*p*_^*2*^*=* 0.131. The fish’s tracking performance was at its peak in stationary (*M* =0.332, *SD* =0.100) compared to other speed levels, p < .001. Moreover low levels (*M* =0.348, *SD* =0.113) and medium levels (*M* =0.348, *SD* =0.133) better than high levels (*M* =0.362, *SD* =0.101) (*p* =0.005, *p* < .001). However, no significant differences were found among the low and medium levels (Fig.5).

**Fig.5.**
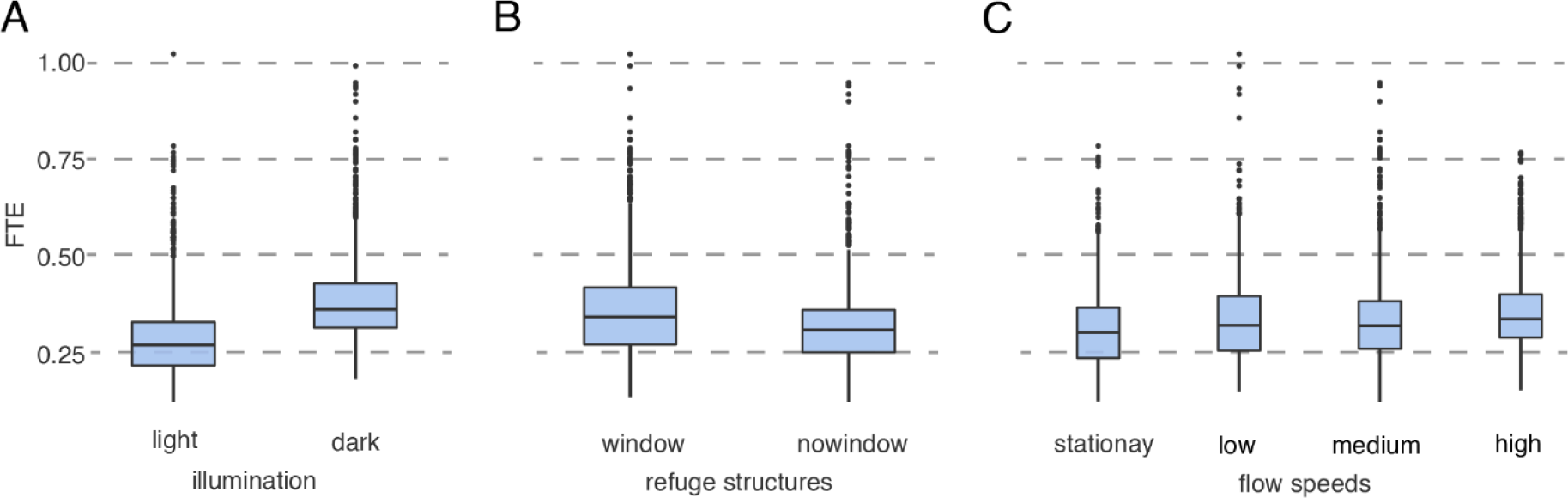
Smooth purusit tracking across various sensory conditions in the frequency domain

**Fig.6.**
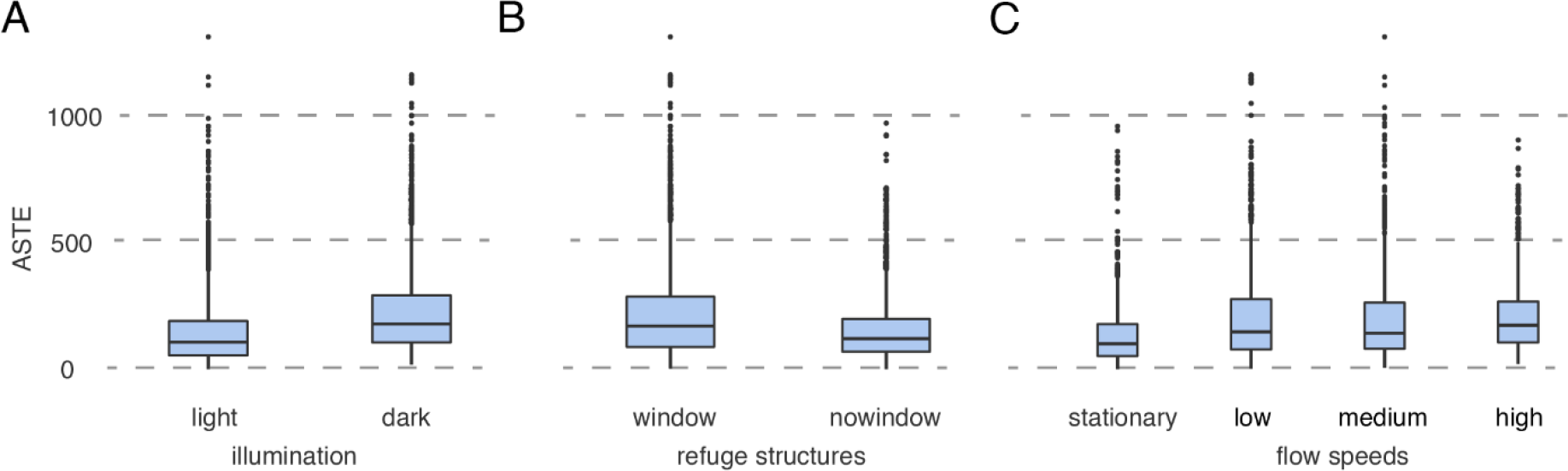
Active sensing movements across various sensory conditions in the frequency domain

Additionnally to these main effects, we identified a significant interaction between illumination conditions and flow speeds, *F* (2.86,684.14) =4.267, *p* =0.006, *h* _*p*_^*2*^*=* 0.018. Also we found a significant interaction between refuge structures and flow speeds, *F* (2.89,689.86) =35.638, *p* < .001, *η*_*p*_^*2*^*=* 0.130. Lastly we found a significant interaction between illumination, refuge structures and flow speeds, *F* (2.87,684.77) =7.790, *p* < .001, *η*_*p*_^*2*^*=* 0.032. Post hoc comparisons with Bonferroni correction revealed that tracking performance was worse in light conditions with windows compared to light conditions without windows (*p* < .001). On the contrary, tracking performance was better in light conditions with windows than in dark conditions with windows. Additionally, tracking performance was better in light conditions without windows than in dark conditions with windows or without windows, with all comparisons being statistically significant (*p* < .001).

### 3.2. The effects of illumination, refuge structure and flow speed on active sensing

All movements other than the frequency of the inputs given here were considered as active sensing movements. These 13 frequencies which are used for smooth pursuit tracking were removed from the fish’s movements and again we used same weighted system for section 3.1 for other active sensing frequencies. Briefly we divided the total result from this weighting by the number of frequencies (237) and defined it as the error corresponding to a single frequency which is active sensing frequency domain tracking error (ASTE).

We detected a significant influence of the illumination environment on tracking performance, showing a substantial difference (*F* (1,239) = 150.9957, *p* < .001, *η*_*p*_^*2*^*=* 0.387). This finding highlighted that tracking performance excelled in light settings (*M* =162, *SD* =145) in contrast to dark conditions (*M* =245, *SD* =180). Our analysis also revealed that the structure of the refuge had a considerable impact on tracking performance (*F* (1,239) = 81.1413, *p* < .001, *η*_*p*_^*2*^*=* 0.253).

To our surprise, contradicting our expectations based on existing literature, tracking performance was superior in situations without windows (*M* = 170, *SD* = 132) compared to those with windows (*M* = 236, *SD* = 193). Furthermore, we observed a primary effect of flow speed on tracking performance (*F* (2.93,700.12) = 65.5279, *p* < .001, *η*_*p*_^*2*^*=* 0.215). The fish’s tracking performance reached its highest point in stationary (*M* = 152, *SD* = 133) compared to other speed levels (low (*M* = 223, *SD* = 196): *p* < .001, medium (*M* = 219, *SD* = 187): *p* < .001 and, high (*M* = 221, *SD* = 139): *p* < .001).

In addition to these primary findings, we observed a substantial interaction between illumination conditions and flow speeds (*F* (2.76,659.24) = 14.2187, *p* = <0.001, *η*_*p*_^*2*^*=* 0.056) and, refuge structures and flow speeds (*F* (2.46,587.16) = 17.5133, *p* < .001, *η*_*p*_^*2*^*=* 0.068).

## 4. DISCUSSION

This research aims to investigate the behavior of weakly electric fish in response to flow speeds similar to what they observe in their natural environment. We study the impact of the flow speed within the context of the refuge tracking behavior of the fish. This tracking behavior, which occurs on a single linear axis, provides a very convenient environment for the investigation of the effect of flow speeds during the unconstrained free swimming behavior of the weakly electric fish.

### 4.1. Flow speeds effect the tracking performance

The reason for determining the flow speed as the aim of this study is that the effect of controlled flow on a free-swimming fish was not examined in the literature before. However, in the direction of this research, we examined the effects of illumination and refuge structures, which were examined before in the literature. Similar to the results presented in the literature, the tracking behavior is better and active sensing movements are less in the light environment than in the dark environment. However, unlike the current results in the literature, a better tracking was observed in the no window case than the case of refuges with windows. We believe that the reason for this difference is that the fish can perceive corners better, since the fish are taller than the height of the refuges.

According to our results, the tracking performance of fish under flow is decreased as it becomes harder for the fish to swim against flow. As the flow speed increased, it became more difficult for the fish to follow the refuge due to the limited thrust force. This caused changes in gain and phase of the tracking response of the fish. The phase gradually increased as the gain gradually decreased. A decrease in gain was expected, because the fish cannot give the desired output to the given input due to a movement against the flow. However, we expected to see an increase in phase lag, since we anticipated that it will take longer for the fish to respond to the given input under flow. However, the results showed that the phase lag was decreased with increasing flow speed. One possible reason for this could be that the fish might be drifting much faster when swimming in the direction of water flow. When the fish swims in opposite direction with the refuge, its controller may compensate for the effects of flow speed. This way, fish might have an advantage in terms of phase lag.

The main reason behind the effects of flow speed is that it modulates the swimming dynamics of the fish. From a modeling perspective, the swimming dynamics, or the plant dynamics, correspond to the mapping from the motor output of the central nervous system and the position of the fish. Changes in the flow speed modulates the mapping from the motor commands to the fish position. Therefore, the effects of flow speed can be consolidated to changes in locomotor dynamics. This also appears in the study conducted by Sefati et al., where different speeds were tested on a robotic weakly electric fish, and in the resulting model, it was seen that the flow speed of the water affected the nodal point, corresponding to the kinematics of the fish (Sefati et al., 2013). Finally, Hawkins et al. and Ortega-Jimenez et al. showed that water flow rate changes the kinematic behavior of fish. These studies looked at the fish’s interactions with the flow and the behaviors used for kinematics, and as a result, it was found that the water flow spped affected the locomotive activities of the fish. However, these studies did not look at how the behavior changes during a goal-oriented task (Hawkins et al., 2022; Ortega-Jiménez and Sanford, 2021).

Finally, the fact that there is no difference in tracking behavior between low speed and medium speed in the frequency domain, but there is a difference of both as compared to high speed. These showed that tracking behavior may get worse as the speed further increases, but we were not able to test in higher speeds as it causes the fish drag with the water. One reason why we observe these differences in the frequency domain but not in the time domain is that we can generated bootstrap copies of the frequency response functions in the frequency domain. This allows working with more data, which emphasizes the differences between the two cases.

### 4.2. Active sensing movements increases under flow

To compensate for the impact of water flow speed, weakly electric fish exhibit active sensing movements. These movements can include adjustments in posture, fin movements, or changes in swimming behavior. By altering their movement patterns, weakly electric fish can enhance their ability to detect and interpret electrical signals in different water flow conditions. These active sensing movements can help them maintain a constant distance from objects of interest, navigate through varied flow conditions, and adapt to changes in their environment.

As a result of our experiments, we observed statistically significant differences between flow speeds and stationary water. We expected this because, as explained in section 4.1, as the water flow speed increases, it will become more difficult for the fish to perform the tracking behavior. Therefore, it will resort to extra movements, namely active sensing movements, to complete the tracking behavior or to continue the movement.

However, no statistical difference was found between low or high flow speeds in terms of the active sensing movements conducted by the fish. The reason for this may be that there is not sufficient difference between flow speeds to trigger a change in the active sensing movements. Another reason may be the decrease in the need for active sensing due to enhanced stimulation of the mechanoreceptors of the fish. The key reason behind the active sensing movements is that fish tries to improves its state estimation performance. It is highly likely that increased mechanosensation contributes to state estimation, and thus reduces the need for active sensing.

In conclusion, the effects of flow speeds in weakly electric fish were investigated in this study. As a result of the research, a significant difference was found between the stationary and the flow speeds. For this reason, this situation should be taken into account in future experiments with these weakly electric fish. Finally, keeping this parameter in mind while modeling the sensor structures of these fish will allow the model to be more realistic.

## Acknowledgements

We thank Noah J. Cowan and Mustafa Mert Ankarali for their invaluable discussions.

## Author contributions

EYA: Co-designed experiments, performed experiments,proceesed and analyzed data, wrote the orginal draft. BU:Analyzed data and review and edit draft. IU: Co-designed experiments, proceesed, analyzed and oversaw data analysis, supervised the project, wrote the orginal draft, funding.

## Data availability

The data and codes will be made available to everyone with doi when the final version of the article is published.

